# Acquirement of the autonomic nervous system modulation evaluated by heart rate variability in medaka (*Oryzias latipes*)

**DOI:** 10.1101/2022.08.02.502546

**Authors:** Tomomi Watanabe-Asaka, Maki Niihori, Kento Igarashi, Shoji Oda, Ken-ichi Iwasaki, Yoshihiko Katada, Toshikazu Yamashita, Hiroki Sonobe, Masahiro Terada, Shoji A. Baba, Hiroshi Mitani, Chiaki Mukai

## Abstract

Small teleosts have recently been established as models of human diseases. However, measuring heart rate by electrocardiography is highly invasive for small fish. The physiological nature and function of vertebrate autonomic nervous system (ANS) modulation of the heart has traditionally been investigated in larvae with an incompletely developed ANS or in anesthetized adults, whose ANS activity may possibly be disturbed under anesthesia. Here, we defined the frequency characteristics of heart rate variability (HRV) modulated by the ANS from observations of heart movement in high-speed movie images and changes in ANS regulation under environmental stimulation in unanesthetized adult medaka (*Oryzias latipes*), a small teleost.

The HRV was significantly reduced by atropine (1 mM) in the 0.25 – 0.65 Hz and by propranolol (100 μM) at 0.65–1.25 Hz range, suggesting that HRV in adult medaka is modulated by both the parasympathetic and sympathetic nervous systems within these frequency ranges. Such modulations of HRV by the ANS were remarkably suppressed in anesthetized adult medaka. Continuous exposure to light suppressed HRV only in the 0.25 – 0.65 Hz range, indicating parasympathetic withdrawal. The power of HRV increased along developmental processes. These results suggest that ANS modulation of the heart in adult medaka is frequency-dependent phenomenon, and that the impact of long-term environmental stimuli on ANS activities can be precisely evaluated in unanesthetized adult fish using this method.

## Introduction

Living organisms continually respond to various types of environmental stress. The autonomic nervous system (ANS) plays an important role in adjusting the various physiological parameters in coordination with the hormonally-regulated endocrine system. The vertebrate heart responds to changes in physical and physiological conditions by regulating the heartbeat and accumulated evidence supports the notion that the vertebrate heart rate is controlled by the ANS, which comprises sympathetic and parasympathetic nervous systems [1]. Both branches of the ANS regulate cardiac activity; the parasympathetic system decreases, whereas the sympathetic system increases steady-state heart rate [2]. The ANS regulates heart rate variability (HRV) in addition to steady-state heart rate. Various types of analysis have been developed to observe HRV regulation by the ANS and to understand how the vertebrate ANS functions in humans and fish [3–7]. Teleost fish are held to be the first in the phylogenetic tree to have a true ganglionated sympathetic trunk together with a distinct vagal system that is similar to that in mammals [8].

The function of the ANS in cardiac regulation has been investigated using electrocardiographic HRV analysis in anesthetized adult fish, as well as heart rate analysis using video imaging in embryos or in larvae just after hatching [9–12]. Analysis using the electrocardiography (ECG) is effective in finding intrabeat abnormalities of the heart, such as QT prolongation. Therefore, studies using large fish such as scorpion fish or rainbow trout have been promoted by implanted electrode [13]. Analysis using an ECG is also used in small fish like zebrafish, however the method using needle electrodes in small fish such as zebrafish requires that placing to expose ventral side and put water directly from the mouth by tube with anesthesia or muscle relaxants. These methods are highly invasive, so that results in serious difficulties with HRV data acquisition from intact fish which is the target of this study [7, 11]. To maximally reduce the impact on the fish, the suppression of ANS activity by anesthesia needs to be eliminated and the invasiveness of the measurements needs to be reduced. However, although the popular anesthetic MS-222 (Tricaine) interferes with sympathovagal function in fish [10, 14], anesthesia is nevertheless required to obtain electrocardiographic measurements from adult fish and amphibians [15–17].

Imaging technologies have options less invasive than electrocardiography. Motion pictures allow analyses of heart development and heartbeat in immobilized embryos and larvae [18–21]. The fish heart comprises a single atrium and a single ventricle, which facilitates optical measurement and analysis. Heart movement in adult small fish can also be measured using infrared light and high-frequency ultrasound, but these methods still require immobilizing adult fish with anesthetics [12, 22].

Small fish such as zebrafish and medaka have recently been established as models of human diseases [23–26]. Several pharmacological studies have recently found that larval-stage zebrafish express receptors for sympathetic and parasympathetic neural transmission and that heart rates in zebrafish larvae change in response to both sympathetic and parasympathetic input with or without anesthesia [16, 20, 27, 28]. However, one of these studies further demonstrated that the autonomic components of the reflex are poorly developed in 5-day-old larval zebrafish, indicating that ANS function is incompletely developed at this stage [20]. Motion pictures are difficult to acquire from highly mobile, opaque adult zebrafish and can only be obtained from transparent immobile embryos or immobilized larvae. Moreover, to evaluate long-term changes in ANS activity such as those induced by environmental changes, adult animals without major alterations in ANS activity during growth or sexual maturation should be studied.

Medaka are small teleosts that are native to East Asian freshwater systems and they have become popular models of human diseases because they are easily maintained in laboratories, their genetics and development are known in detail, and their whole genome has been sequenced [29, 30]. In addition, medaka have several characteristic features that facilitates the study of ANS activities in unanesthetized adults: they are highly adaptive to a wide range of temperatures as well as to low oxygen content, prefer slowly flowing water and do not swim vigorously. The transparent strain, SukeSuke (SK2), has been established by crossing several spontaneous body-coloring mutants [31, 32] and SK2 heart movement can be observed through their transparent peritoneum. The present study determines the frequency characteristics of HRV modulation by the ANS and non-invasively quantifies ANS activities from spectral analyses of HRV in unanesthetized medaka using high-speed movie images of the heart.

## Materials and Methods

### Medaka strains and husbandry

Specimens of the medaka (*Oryzias latipes*) strain SukeSuke (SK2) reared in common tanks was obtained from our breeding colony. All experiments were proceeded on adult fish over 3 months after hatching whose body lengths were 2.5 ± 0.2 cm. We also analyzed embryos at 6 days post-fertilization at the stage of heart function development (embryonic stage (St.) 36) [33]. The SK2 strain is homozygous for three recessive pigmentation mutations (*b^g8^*; null melanophore, *lf*; leucophore free, *gu*; guanineless) [31, 32] and without apparent abnormalities in cardiac activity during embryogenesis or as an adult. The fish and embryos were maintained under standard laboratory conditions at 26°C with a 14:10-h light-dark cycle in an incubator at 26°C. We maintained fish in tanks under light for 24 h/day for one week before HRV measurements (continuous light). All experiments were performed between 15:00 – 17:00 to avoid diurnal variations. Committees for Institutional Animal Care of the Japan Aerospace Exploration Agency and of the University of Tokyo approved the animal protocols.

### Pharmacology and reagents

Atropine (1 mM), propranolol (100 μM) or MS-222 (80 μg/mL Tricaine, Sigma-Aldrich) were added to a bath containing a small tank holding the fish [14]. Fish were acclimated to the observation container for 5 min before assays. Atropine or propranolol was administered to fish 5 min after starting the assays. MS-222 was administered for a minimum of 5 min before assays. Images of the heart area were taken for 20 min throughout the assays.

### Imaging system and digital video recording

Digital video recording of cardiac activities of adult fish and hatchlings were prepared as described before [34]. Water and oxygen and maintained at 25 ± 1 °C throughout heart movement recording. The swimming area for the fish was partially restricted without affecting ventilation and the heart rate of the fish in the tank was not altered for any longer than two hours. Each measurement was completed within 20 min, including the time to place the embryos into tanks and to establish a quiet, resting and steady state (usually within about 5 min). Heart movement in adult medaka was recorded by taking video movies of the ventral view of adult SK2 through the transparent peritoneum using an inverted stereomicroscope (LEICA Fluorescent Dissecting Microscope MZFLIII with a PLANAPO 1.0X lens) equipped with a digital high-speed camera (CASIO Exilim EX-F1) at 2.0X magnification [35].

Dechorionated embryos at St. 36 [33] were immobilized in the orientation appropriate for video recording using 2.5% methylcellulose in glass-bottomed dishes (Matsunami, Japan) and then heart movement was recorded using a stereomicroscope equipped with a digital high-speed camera (EX-F1) as described before [36]. Digital pictures at 300 frames per second (fps) with a resolution of 512 x 384 pixels were captured for up to 20 min and recorded in a PC using Final Cut Pro software (Apple Computer). Since 2 – 5 min data acquisition is recommended for HRV analysis in humans [4] and heart rate is faster in medaka than in humans, we acquired data for 3 minutes after recovery from heart rate acceleration caused by being placed in the tank. We defined a stable heart rate as steady-state, which was similar (± 10 bpm) between the first and last 3 min.

### Extraction of cardiac activity

We extracted the cardiac activities of adult fish, juveniles, larvae and embryos according to the previous papers [34, 36]. The pixel intensities of each ROI were digitalized throughout the entire time series examined using Bohboh software (Bohboh Soft, Tokyo, Japan) and further processed using Cutwin mathematical software (EverGreen Soft, Tokyo, Japan). Data were processed by taking a moving average over 21 frames. Finally, local maxima and minima detected using Cutwin software were identified as the end of systole and of diastole, respectively. The pixel intensity of the ROIs in the heart images of immobilized embryos was digitalized, movement-averaged over 21 frames and then local maxima and minima were similarly determined as described above.

### Steady state heart rate, respiratory rate and power spectral analysis of HRV

The period between pixel intensity minima representing the end of diastole provided the interbeat interval from which we calculated beat-by-beat heart rate. We then averaged beat-by-beat heart rates during collection for 3 min to generate steady-state heart rates. Respiratory rates per minute were determined by counting the number of opercula movements in 30 sec of data collection for 3 min. Beat-by-beat heart rates were linearly interpolated and resampled at 8 Hz to create an equidistant time series for spectral analyses of HRV. The time series of heart rates was initially detrended with third-order polynomial fitting and then subdivided into 512-point segments with a 50% overlap, resulting in five data segments collected over a period of 3 min. Fast Fourier transform (FFT) was applied to obtain a power spectrum of HRV on each Hanning-windowed data segment and subsequently the power spectra of the five segments were subsequently averaged to calculate the autospectrum of HRV acquired during 3 min. The minimal resolution of these spectra was 0.015625 Hz. The data described above were processed using DADiSP software (DSP Development, Cambridge, MA, USA).

### Statistical analyses

Data were statistically analyzed using a one-way ANOVA followed by a comparison with control (Dunnett’s post hoc test) using JMP software (SAS Japan, Tokyo, Japan). A P value of < 0.05 was considered statistically significant. Data are presented as means ± s.d. of five fish or embryos per experiment.

## Results

### Steady-state heart rate and respiratory rate

The steady-state heart rate in the control fish was 137.1 ± 6.70 bpm (Fig 1A, n = 5). Atropine increased the rate to 164.8 ± 9.69 bpm (n = 5, *p* = 0.009), showing that atropine induced tachycardia compared with the control, whereas propranolol induced bradycardia by decreasing the heart rate to 106.3 ± 14.6 bpm (Fig 1A, n = 5, *p* = 0.047). The steady-state heart rates in adult fish under anesthesia with 80 μg/mL of MS-222 and under continuous light conditions, were 149.6 ± 12.0 (n = 5) and 147.9 ± 9.07 (n = 5) bpm, respectively (Fig 1A), which did not significantly differ from those of the control (*p* = 0.20 and 0.37, respectively).

**Fig 1.**
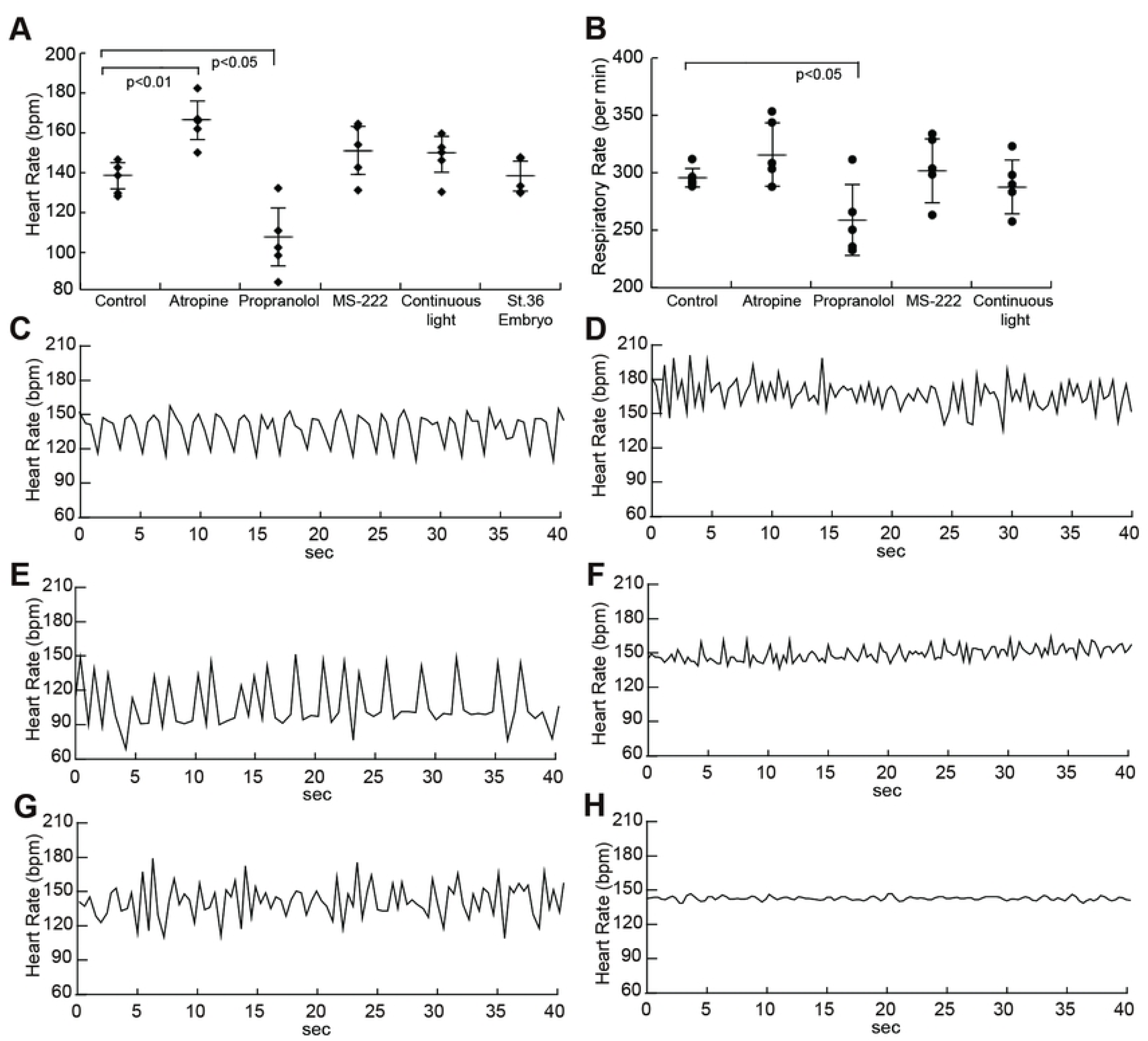
Average of steady-state heart rate, respiratory rate and examples of beat-by-beat heart rates. Group averaged steady-state heart rate were plotted under each condition (A). Group averaged respiratory rate (per min) were plotted under each condition in adult medaka (B). Standard deviations among five fish are plotted as error bars. Examples of beat-by-beat heart rate changes for 40 s under each experimental condition are presented (C-H). Control (C), 1 mM atropine (D), 100 μM propranolol (E), anesthesia with 80 μg/mL MS-222 (F), continuous light (G) and St. 36 embryo (H).

The respiratory rate in the control fish was 296 ± 8.22 per min (Fig 1B) and the rates in adult fish administered with propranolol were 259 ± 31.1 per min (Fig 1B), and significantly decrease from control values (*p* = 0.033). The rates in adult fish administered with atropine and MS-222 and adult fish under constant light were 316 ± 27.7, 302 ± 28.0 and 288 ± 23.6 per min, respectively (Fig 1B), which did not significantly differ from control values.

### Control HRV

Fig 1C shows an example of heart rate over a 40-sec period within a 3-min sample from a control fish. Specific rhythms seemed to emerge in the form of definite heart rate fluctuations. The mean of the power spectral density of the HRV in five adult intact fish (control) shows oscillatory periods at frequencies below 1.25 Hz, and at least two specific peaks (Fig 2A, black line). Power spectral density at a frequency of 0 Hz was omitted from this analysis and the power spectrum was divided into low- (0.02–0.25 Hz, Fig 2B), middle- (0.25–0.65 Hz, Fig 2C) and high- (0.65–1.25 Hz, Fig 2D) frequency ranges. The power of these ranges in the control fish was 48.2 ± 24.8, 171 ± 115 and 98.9 ± 40.3 bpm^2^, respectively.

**Fig 2.**
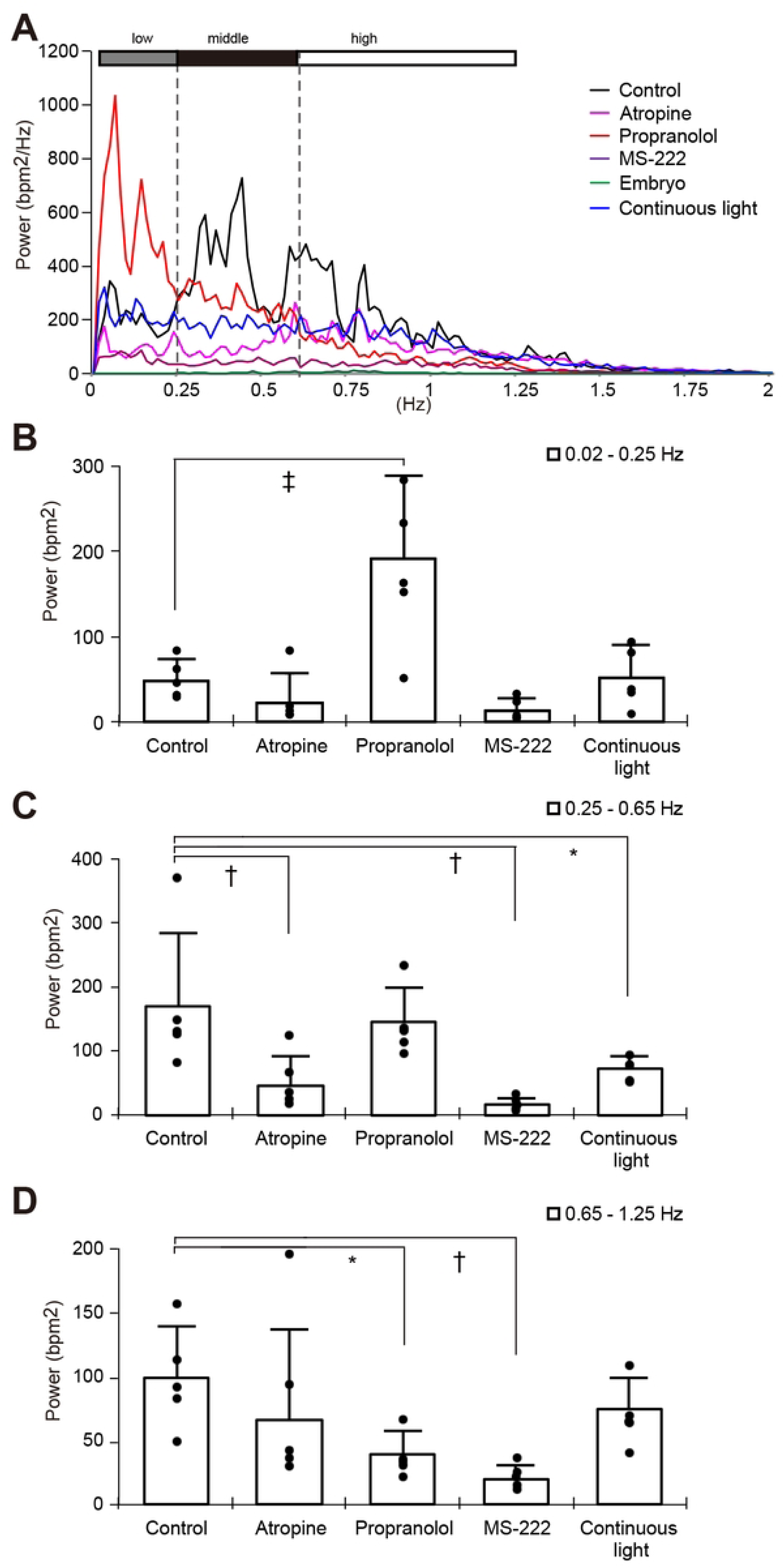
Power spectral analysis of HRV for 3 min in medaka. Power spectral density of HRV in adult control fish, administered with 1 mM atropine, 100 μM propranolol, 80 μg/mL MS-222 and under 1-week continuous light and St. 36 embryos was obtained by fast Fourier transformation using DADiSP software. The mean of power spectral density of HRV in five fish under each condition is shown. Low (0.015625–0.25 Hz), middle (0.25–0.65 Hz) and high (0.65–1.25 Hz) frequencies are shown as gray, black and white bars, respectively (A). The amount of power was added in each frequency range and error bars are standard deviations among five fish. Data from individual fish are indicated as black dots (B–D). B, 0.01625–0.25 Hz. C, 0.25–0.65 Hz. D, 0.65–1.25 Hz. \**P* < 0.05, ^†^*P* < 0.01, ^‡^*P* < 0.005, compared with control.

### Effect of inhibitors for autonomic nervous system on HRV

Figs 1D – F show the heart rate fluctuation induced by 1 mM atropine, 100 μM propranolol and 80 μg/mL of MS-222, respectively. The mean of the power of the HRV in five adult fish that were administered with atropine, propranolol and MS-222 of each was shown in Fig 2A (n = 5). The mean power of the low-, middle- and high-frequency ranges in the presence of atropine was 22.8 ± 34.3, 48.0 ± 44.9 and 66.2 ± 70.9 bpm^2^, respectively. The power of the low-, middle- and high-frequency ranges with propranolol administration was 191 ± 96.2, 147 ± 54.4 and 39.4 ± 17.8 bpm^2^, respectively. And those in the low-, middle- and high-frequency ranges under MS-222 anesthesia was 13.9 ±13.8, 18.0 ± 9.40 and 20.3 ± 10.1 bpm^2^, respectively. Atropine significantly reduced the power in the middle-frequency range but not with propranolol (Fig 2C). On the other hand, propranolol increased and decreased the power in the low- and high-frequency ranges despite atropine did not affect in those ranges (Figs 2B and D, respectively). Reductions in the powers of the middle- and high- frequency ranges were statistically significant in MS-222 administered fish (Figs 2B – D).

### HRV under continuous light

Fig 1G shows heart rate fluctuation in adults after one week under continuous light. The power of the low-, middle- and high-frequency ranges of these fish was 51.7 ± 38.2, 75.2 ± 18.7 and 75.0 ± 24.6 bpm^2^, respectively (Fig 2A blue line; Figs 2B – D). The power in the middle-frequency band was significantly reduced under continuous light for 1 week (Fig 2C).

### HRV acquirement in medaka

Figs 3A and B show local minima and maxima identified as the end of diastole and of systole, respectively. Fig 3C shows changes in intensity inside the white ROI in the heart area as an indicator of heartbeat for 20 sec (Figs 3A and B; white circles). Thus, heartbeats were clearly determined from changes in pixel intensity of the heart area in St. 36 medaka embryos.

**Fig 3.**
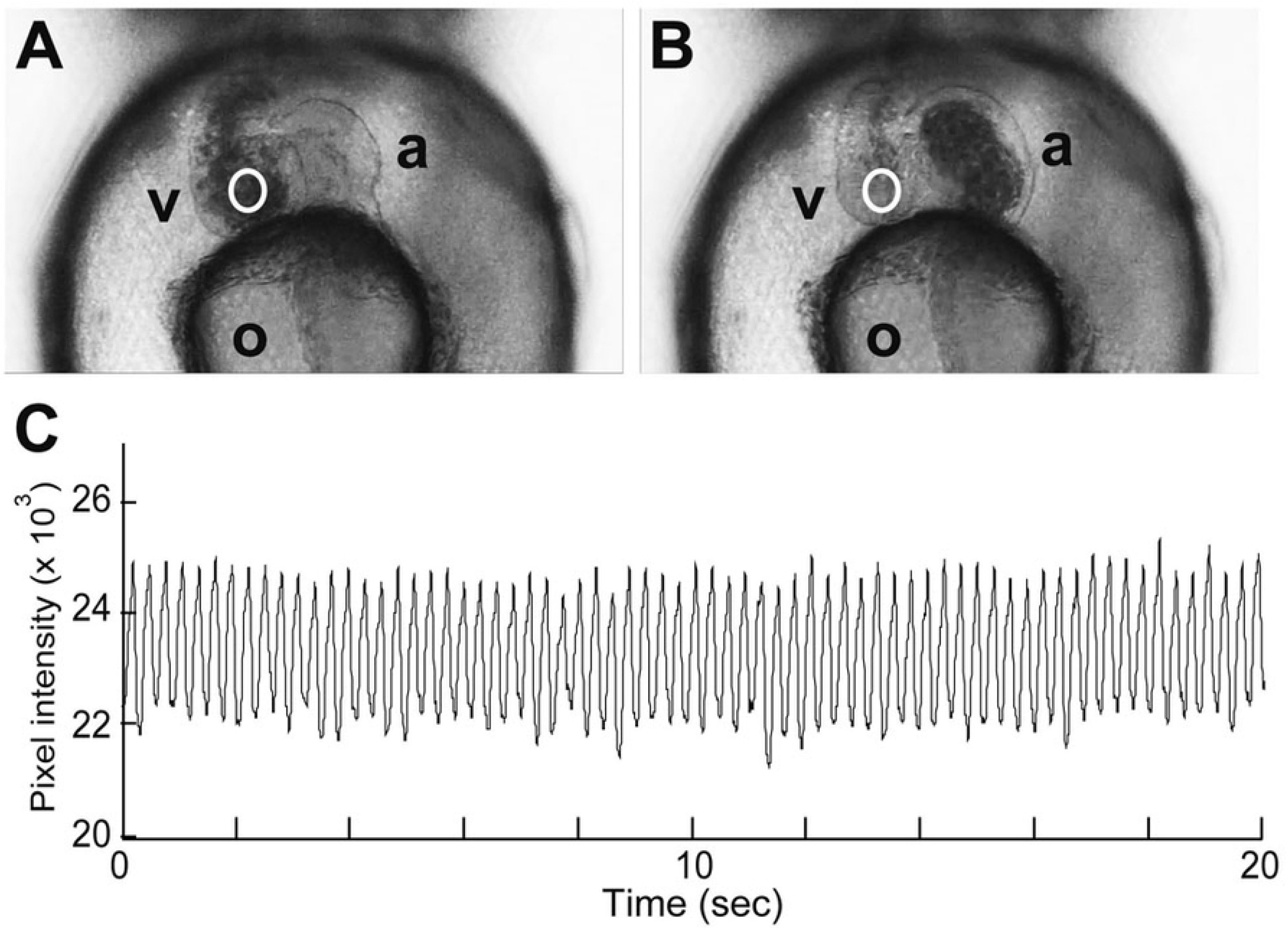
Automated extraction of heart movement in embryos. Sample pictures are at diastole and systole (A and B) in embryonic heart area view. Pixel intensities in the white circular ROI were extracted as numerical data. Twenty seconds of total intensity in white circles throughout the entire series were plotted after being smoothed by the 21-point moving average (C). a, atrium; v, ventricle.

**Fig 4.**
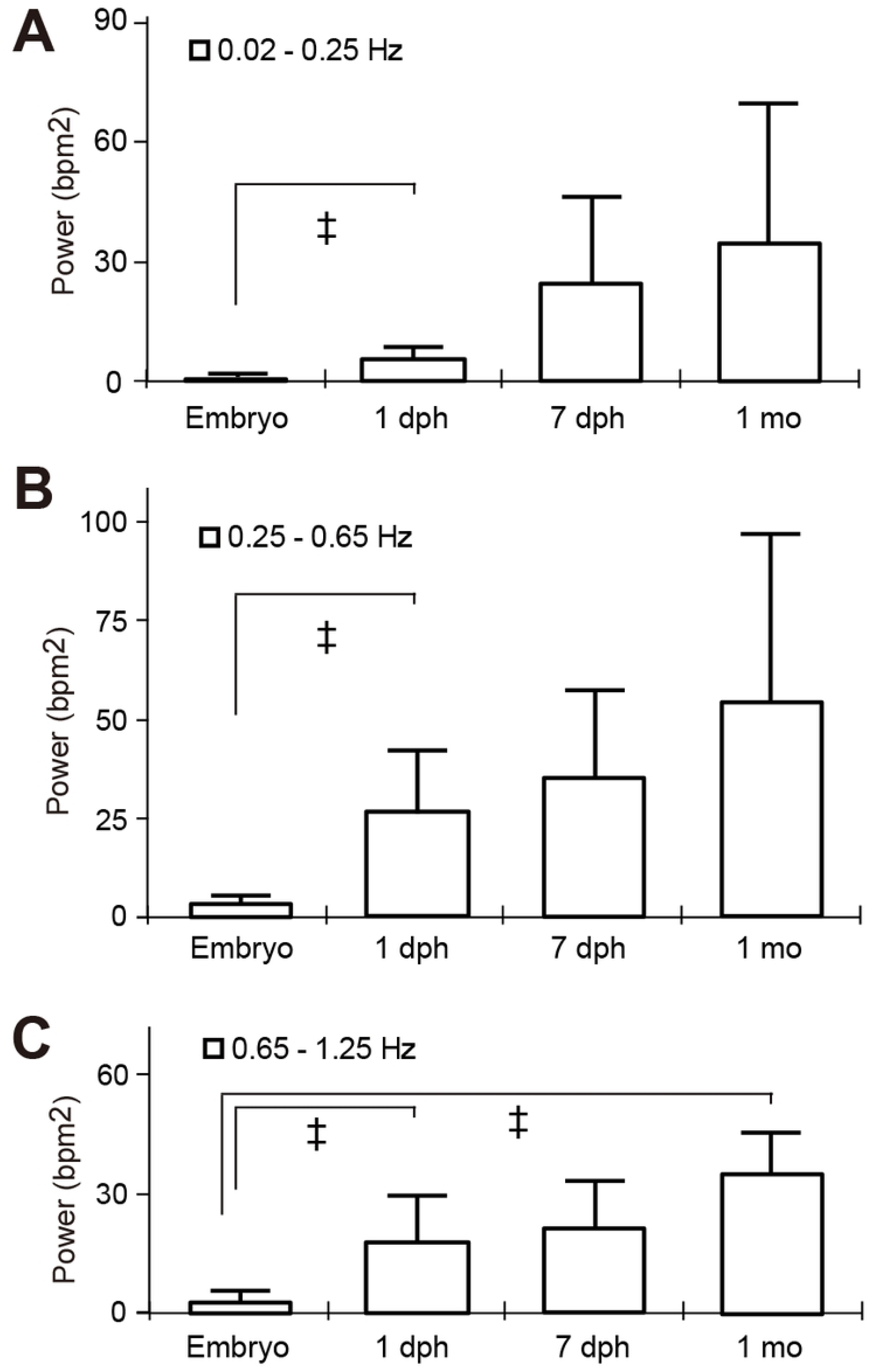
Power spectral analysis of HRV for 3 min according to development. Power spectral density and the mean of power spectral density of HRV in st.36 embryos (n = 5), 1 day post hatch (dph) larvae (n = 20), 7 dph larvae (n = 3) and 1 month old juvenile (n = 4) was obtained by fast Fourier transformation using DADiSP software. Low (0.015625–0.25 Hz), middle (0.25–0.65 Hz) and high (0.65–1.25 Hz) frequencies are shown in panels A – C, respectively. The amount of power was added in each frequency range and error bars are standard deviations among samples. A, 0.01625–0.25 Hz. B, 0.25–0.65 Hz. C, 0.65–1.25 Hz. ^‡^*P* < 0.005; compared with st.36 embryos.

The beat-by-beat heart rate in St. 36 embryos essentially remained consistent with minimal fluctuation (Fig 1H). To evaluate the development of the ANS modulation during fish grow, we took the high-speed movies for cardiac activities and analyzed the HRV in st. 36 embryos (n = 5), 1 day post hatch (dph, n = 20) and 7 dph larvae (n = 3) and 1 month old juvenile fish (n = 4). The mean power of the low frequency ranges in embryos, 1 dph, 7 dph and 1 month old fish was 0.32 ± 0.31, 5.43 ± 2.91 (*p* < 0.001), 24.4 ± 21.7 (*p* = 0.101) and 34.6 ± 34.9 (*p* = 0.104) bpm^2^, respectively. The mean power of the middle frequency ranges in each stage fish was 3.39 ± 2.11, 26.3 ± 15.7 (*p* < 0.001), 35.0 ± 22.0 (*p* = 0.063) and 53.7 ± 41.0 (*p* = 0.677) bpm^2^, respectively. The mean power of the high frequency ranges in each developmental stage was 3.05 ± 2.08, 18.0 ± 11.8 (*p* < 0.001), 21.2 ± 11.6 (*p* = 0.054) and 35.3 ± 10.8 (*p* = 0.006) bpm^2^, respectively. There was a tendency of increasing the ANS modulation in all frequency ranges associate with the fish development.

## Discussion

We measured steady-state heart rate and HRV in intact adult medaka by extracting heart motion data from ventral high-speed video images and spectral analysis. We then defined the characteristics of the HRV regulation by the parasympathetic and sympathetic nervous systems in adult medaka based on these findings.

### Steady-state heart rate measurement and HRV analysis in medaka

Previous studies of steady-state heart rates in adult medaka have mainly focused on the effects of temperature on the heartbeat of the isolated heart or on the heart of intact adult medaka. These studies found that the steady-state heart rate of medaka is about 140 bpm at 25°C with or without anesthesia [37–42], with which our findings are consistent. Despite interest in comparative studies of cardiac regulation in vertebrates, only a few investigators have applied spectral analysis to non-mammalian vertebrates. The wide diversity of cardiac-related signals, non-standardized procedures and techniques might have hindered the application of spectral analysis to fish [10]. Although several studies have examined ANS involvement in heart rate regulation, variations in specific frequency ranges of the HRV in fish have not been quantified.

We found here that specific peaks appear in the HRV spectrum of control adult medaka and that the power spectrum of HRV in these fish covers at least three frequency ranges (0.02–0.25, 0.25–0.65 and 0.65–1.25 Hz) presumably because of the regulatory machineries discussed below.

### Contribution of the ANS to HRV

Atropine, a muscarinic receptor antagonist, reduced fluctuations in the range of 0.25–0.65 Hz (middle-frequency range) and induced tachycardia. Propranolol, a β-adrenergic receptor antagonist, reduced fluctuation in the range of 0.65–1.25 Hz and induced bradycardia. These results suggested that the parasympathetic and sympathetic nervous system primarily modulate HRV in the middle- and high-frequency ranges, respectively. Anesthesia with MS-222 suppressed fluctuations within both the 0.65–1.25 Hz and 0.25–0.65 Hz ranges and confirmed the findings, that local anesthetics block both the sympathetic and parasympathetic nervous systems [43].

Such correspondence of the sympathetic or parasympathetic nervous system with two frequency bands in medaka seems to contradict the mammalian system. The central circuit in the mammalian sympathetic nervous system is thought to have become slower as network complexity has increased due to an evolutionary increase in the number of synapses [44, 45]. Teleost fish are located nearer the origin than mammals in phylogenetic trees having a true ganglionated sympathetic trunk and a distinct vagal system similar to that of mammals, indicating that these fish have a primitive sympathetic circuit. Moreover, respiration is slower than heart rate in mammals, in which the high frequency range of HRV refers to vagal nerve activity that mainly reflects respiratory sinus arrhythmia. However, ventilation caused by opercula movement is faster than the heart rate in fish and the reported frequency range of such movement in medaka is 4–5 Hz at 25°C [37]. Therefore, the effect of respiratory sinus arrhythmia seen in mammals on HRV would be quite small in teleost fish. Evolutionary aspects and respiration style can explain the difference in autonomic modulation of the frequency bands between medaka and mammals.

Although both atropine and anesthesia by MS-222 tended to suppress fluctuations in the low-frequency range, the difference did not reach statistical significance and the fluctuation in this range considerably differed among individual adult fish. Moreover, propranolol remarkably increased fluctuations in the low-frequency range. Bradycardia and occasional rapid movement developed in adult medaka administered with propranolol, presumably because blood flow was decreased, which increased power in the low frequency range. Such increased power could also be attributed to ventricular arrhythmia caused by the induced bradycardia. Therefore, whether the power in the low-frequency range induced in this manner can serve as an effective indicator of ANS activities requires some consideration. Further studies should examine HRV especially at this frequency range using electrocardiography.

### Influence of environmental disturbance on cardiac ANS

We evaluated the effects of an environmental disturbance caused by exposure to constant light for one week on the ANS activity in adult medaka. The results showed that only HRV decreased only in the middle-frequency range. Since fluctuations in the middle- or high-frequency ranges could indicate parasympathetic or sympathetic nervous activity, respectively, the data suggest that only parasympathetic nervous activity was reduced under continuous light in adult medaka, whereas sympathetic nervous activity was less affected.

### Development of the ANS modulation in medaka

In this study, HRV was hardly observed in st.36 medaka embryos which was before hatching, but there was a tendency of increase in accordance with the growth in one month after hatching. Cardiac branches of the autonomic nerve have also been observed in medaka embryos before hatching [36]. These results suggested that to acquire sufficient function for the autonomic nervous system will take at least one month time after being formed as a structure.

### Advantages of using adult medaka as a model animal

Our findings showed that environmental stimuli and anesthesia can both alter the power spectrum and that ANS activity can be evaluated in unanesthetized adult medaka, when HRV is regulated by both the sympathetic and parasympathetic nervous systems. We found a constant heart rate in medaka embryos, indicating that ANS modulation of HRV has not yet developed in St. 36 medaka embryos, unlike in zebrafish larvae. The use of unanesthetized adult medaka offers advantages for evaluating the impact of environmental stress on ANS activities.

Small fish such as medaka can be reared under the same experimental conditions from the egg to adulthood, and experimental conditions including breeding temperature, lighting schedule and feeding conditions can be easily controlled. Although small fish are difficult to manipulate and cardiac activity is difficult to analyze, small teleosts bred in laboratories are advantageous for studying the effects of environmental conditions on vertebrate ANS activity and development. The transparent adult medaka used herein enabled ANS studies without anesthetics. The medaka system also confers advantages for drug screening and phenotypic analysis of spontaneous or genetically manipulated mutants.

### Study limitations

A type II error in the present study is possible owing to the low sample size. Although HRV in the 0.25–0.65 Hz range significantly decreased in the anesthetized fish and in those under constant light conditions compared with control fish, the steady-state heart rate did not differ from controls despite tendencies towards tachycardia. However, this might also indicate that HRV spectral indices can detect differences in ANS modulation more sensitively than steady-state heart rate. The resampling process might also have acted as a low-pass filter inducing attenuation at the high frequency range.

The number of the opercula movements did not significantly differ, although they tended to increase and decrease under the administration of atropine and propranolol, respectively. Fish exchange gases through branchial respiration and more energy is required for ventilation with gills than with pulmonary respiration [46]. Therefore, HRV regulation by respiratory rate is likely to be more complex in fish than in mammals, despite a correlation between heart rate and respiratory rates [47]. Since we excluded the effects of ventilation movements by data processing before spectral analysis and the number of the opercula movements did not significantly differ, we might have minimized the influence of ventilation frequency in HRV.

In summary, we identified the steady-state heart rate in medaka by extracting heart motion from ventral images and applying spectral analysis, and characterized part of the frequency nature of HRV modulation. Atropine significantly reduced HRV in the middle-frequency range of adult medaka, suggesting primarily parasympathetic nervous regulation of HRV within this range. Propranolol significantly reduced HRV in the high-frequency range of adult medaka, suggesting sympathetic nervous regulation within this frequency range. Such HRV modulations were assessed from embryo to adult fish in the same system by using this method. Moreover, constant light reduced HRV only at the middle-frequency range, suggesting the induction of suppressed parasympathetic nervous activity. The present findings constitute a major contribution to comprehending precise ANS modulations caused by environmental stimuli and the processes of ANS maturation.

## Acknowledgements

We are obliged to Dr. Hiroshi Ohshima and Mr. Masafumi Yamamoto for laboratory management. We also express our appreciation to Drs. Tomoaki Matsuo, Masamichi Sudoh, Satomi Tanimoto and Toshiko Ohta for helpful discussions and constructive advice.

## Funding

This study was part of the JAXA-ISS Space Biomedical Research Project (2009–2012) at the Japan Aerospace Exploration Agency and supported by the Ministry of Education, Culture, Sports, Science and Technology [KAKENHI 24310039 to S.O.].

## Disclosure statement

The authors have nothing to declare.

## Notes

### Competing Interest Statement

The authors have declared no competing interest.

## References

1. Levy MN. Sympathetic-parasympathetic interactions in the heart. Circ Res. 1971 Nov;29(5):437–445. https://doi.org/10.1161/01.res.29.5.437 PMID: 4330524.

2. Steele SL, Yang X, Debiais-Thibaud M, Schwerte T, Pelster B, Ekker M, et al. In vivo and in vitro assessment of cardiac beta-adrenergic receptors in larval zebrafish (Danio rerio). J Exp Biol. 2011 May 1;214(Pt 9):1445–1457. https://doi.org/10.1242/jeb.052803 PMID: 21490253.

3. Akselrod S, Gordon D, Ubel FA, Shannon DC, Berger AC, Cohen RJ. Power spectrum analysis of heart rate fluctuation: a quantitative probe of beat-to-beat cardiovascular control. Science. 1981 Jul 10;213(4504):220–222. https://doi.org/10.1126/science.6166045 PMID: 6166045.

4. Heart rate variability. Standards of measurement, physiological interpretation, and clinical use. Task Force of the European Society of Cardiology and the North American Society of Pacing and Electrophysiology. Eur Heart J. 1996 Mar;17(3):354–381. PMID: 8737210.

5. Sun P, Zhang Y, Yu F, Parks E, Lyman A, Wu Q, et al. Micro-electrocardiograms to study post-ventricular amputation of zebrafish heart. Ann Biomed Eng. 2009 May;37(5):890–901. https://doi.org/10.1007/s10439-009-9668-3 PMID: 19280341; PMCID: PMC6991467.

6. Taylor EW, Leite CA, Skovgaard N. Autonomic control of cardiorespiratory interactions in fish, amphibians and reptiles. Braz J Med Biol Res. 2010 Jul;43(7):600–610. https://doi.org/10.1590/s0100-879x2010007500044 PMID: 20464342.

7. Yu F, Li R, Parks E, Takabe W, Hsiai TK. Electrocardiogram signals to assess zebrafish heart regeneration: implication of long QT intervals. Ann Biomed Eng. 2010 Jul;38(7):2346–2357. https://doi.org/10.1007/s10439-010-9993-6 PMID: 20221900; PMCID: PMC3117900.

8. Nilsson S. Autonomic nerve function in the vertebrates (13). Springer Science & Business Media. 2012.

9. Altimiras J. Understanding autonomic sympathovagal balance from short-term heart rate variations. Are we analyzing noise? Comp Biochem Physiol A Mol Integr Physiol. 1999 Dec;124(4):447–460. https://doi.org/10.1016/s1095-6433(99)00137-3 PMID: 10682243.

10. Campbell HA, Klepacki JZ, Egginton S. A new method in applying power spectral statistics to examine cardio-respiratory interactions in fish. J Theor Biol. 2006 Jul 21;241(2):410–419. https://doi.org/10.1016/j.jtbi.2005.12.005 PMID: 16443239.

11. Campbell HA, Taylor EW, Egginton S. The use of power spectral analysis to determine cardiorespiratory control in the short-horned sculpin Myoxocephalus scorpius. J Exp Biol. 2004 May;207(Pt 11):1969–1976. https://doi.org/10.1242/jeb.00972 PMID: 15107449.

12. Sun L, Lien CL, Xu X, Shung KK. In vivo cardiac imaging of adult zebrafish using high frequency ultrasound (45-75 MHz). Ultrasound Med Biol. 2008 Jan;34(1):31–39. https://doi.org/10.1016/j.ultrasmedbio.2007.07.002 PMID: 17825980; PMCID: PMC2292109.

13. Milan DJ, Jones IL, Ellinor PT, MacRae CA. In vivo recording of adult zebrafish electrocardiogram and assessment of drug-induced QT prolongation. Am J Physiol Heart Circ Physiol. 2006 Jul;291(1):H269–H273. https://doi.org/10.1152/ajpheart.00960.2005 PMID: 16489111.

14. Rombough PJ. Ontogenetic changes in the toxicity and efficacy of the anaesthetic MS222 (tricaine methanesulfonate) in zebrafish (Danio rerio) larvae. Comp Biochem Physiol A Mol Integr Physiol. 2007 Oct;148(2):463–469. https://doi.org/10.1016/j.cbpa.2007.06.415 PMID: 17643329.

15. Hedrick MS, Winmill RE. Excitatory and inhibitory effects of tricaine (MS-222) on fictive breathing in isolated bullfrog brain stem. Am J Physiol Regul Integr Comp Physiol. 2003 Feb;284(2):R405–R412. https://doi.org/10.1152/ajpregu.00418.2002 PMID: 12414435.

16. Huang WC, Hsieh YS, Chen IH, Wang CH, Chang HW, Yang CC, et al. Combined use of MS-222 (tricaine) and isoflurane extends anesthesia time and minimizes cardiac rhythm side effects in adult zebrafish. Zebrafish. 2010 Sep;7(3):297–304. https://doi.org/10.1089/zeb.2010.0653 PMID: 20807039.

17. Randall DJ. Effect of an anaesthetic on the heart and respiration of teleost fish. Nature. 1962 Aug 4;195:506. https://doi.org/10.1038/195506a0 PMID: 14490222.

18. Chan PK, Lin CC, Cheng SH. Noninvasive technique for measurement of heartbeat regularity in zebrafish (Danio rerio) embryos. BMC Biotechnol. 2009 Feb 19;9:11. https://doi.org/10.1186/1472-6750-9-11 PMID: 19228382; PMCID: PMC2664803.

19. Fink M, Callol-Massot C, Chu A, Ruiz-Lozano P, Izpisua Belmonte JC, Giles W, et al. A new method for detection and quantification of heartbeat parameters in Drosophila, zebrafish, and embryonic mouse hearts. Biotechniques. 2009 Feb;46(2):101–113. https://doi.org/10.2144/000113078. PMID: 19317655; PMCID: PMC2855226.

20. Mann KD, Hoyt C, Feldman S, Blunt L, Raymond A, Page-McCaw PS. Cardiac response to startle stimuli in larval zebrafish: sympathetic and parasympathetic components. Am J Physiol Regul Integr Comp Physiol. 2010 May;298(5):R1288–1297. https://doi.org/10.1152/ajpregu.00302.2009 Epub 2010 Feb 3. PMID: 20130228.

21. Taylor EW, Leite CA, Levings JJ. Central control of cardiorespiratory interactions in fish. Acta Histochem. 2009;111(3):257–267. https://doi.org/10.1016/j.acthis.2008.11.006 PMID: 19193400.

22. Yoshida M, Hirano R, Shima T. Photocardiography: a novel method for monitoring cardiac activity in fish. Zoolog Sci. 2009 May;26(5):356–61. https://doi.org/10.2108/zsj.26.356 PMID: 19715506.

23. Martin JS, Renshaw SA. Using in vivo zebrafish models to understand the biochemical basis of neutrophilic respiratory disease. Biochem Soc Trans. 2009 Aug;37(Pt 4):830–837. https://doi.org/10.1042/BST0370830 PMID: 19614603.

24. Matsui H, Ito H, Taniguchi Y, Inoue H, Takeda S, Takahashi R. Proteasome inhibition in medaka brain induces the features of Parkinson’s disease. J Neurochem. 2010 Oct;115(1):178–187. https://doi.org/10.1111/j.1471-4159.2010.06918.x PMID: 20649841.

25. Matsumoto T, Terai S, Oishi T, Kuwashiro S, Fujisawa K, Yamamoto N, et al. Medaka as a model for human nonalcoholic steatohepatitis. Dis Model Mech. 2010 Jul-Aug;3(7-8):431–40. https://doi.org/10.1242/dmm.002311 PMID: 20371730.

26. Watanabe-Asaka T, Mukai C, Mitani H Technologies and Analyses Using Medaka to Evaluate Effects of Space on Health. Biol Sci Space. 2010;24(1):3–9. https://doi.org/10.2187/bss.24.3

27. Jacob E, Drexel M, Schwerte T, Pelster B. Influence of hypoxia and of hypoxemia on the development of cardiac activity in zebrafish larvae. Am J Physiol Regul Integr Comp Physiol. 2002 Oct;283(4):R911–7. https://doi.org/10.1152/ajpregu.00673.2001 PMID: 12228061.

28. Schwerte T, Prem C, Mairösl A, Pelster B. Development of the sympatho-vagal balance in the cardiovascular system in zebrafish (Danio rerio) characterized by power spectrum and classical signal analysis. J Exp Biol. 2006 Mar;209(Pt 6):1093–100. https://doi.org/10.1242/jeb.02117 PMID: 16513936.

29. Kasahara M, Naruse K, Sasaki S, Nakatani Y, Qu W, Ahsan B, et al. The medaka draft genome and insights into vertebrate genome evolution. Nature. 2007 Jun 7;447(7145):714–9. https://doi.org/10.1038/nature05846 PMID: 17554307.

30. Wittbrodt J, Shima A, Schartl M. Medaka--a model organism from the far East. Nat Rev Genet. 2002 Jan;3(1):53–64. https://doi.org/10.1038/nrg704.PMID: 11823791.

31. Fukamachi S, Kinoshita M, Tsujimura T, Shimada A, Oda S, Shima A, et al. Rescue from oculocutaneous albinism type 4 using medaka slc45a2 cDNA driven by its own promoter. Genetics. 2008 Feb;178(2):761–9. https://doi.org/10.1534/genetics.107.073387 PMID: 18245373; PMCID: PMC2248340.

32. Shimada A, Fukamachi S, Wakamatsu Y, Ozato K, Shima A. Induction and characterization of mutations at the b locus of the medaka, Oryzias latipes. Zoolog Sci. 2002 Apr;19(4):411–7. https://doi.org/10.2108/zsj.19.411 PMID: 12130818.

33. Iwamatsu T. Stages of normal development in the medaka Oryzias latipes. Mech Dev. 2004 Jul;121(7-8):605–18. https://doi.org/10.1016/j.mod.2004.03.012 PMID: 15210170.

34. Nagata K, Hashimoto C, Watanabe-Asaka T, Itoh K, Yasuda T, Ohta K, et al. In vivo 3D analysis of systemic effects after local heavy-ion beam irradiation in an animal model. Sci Rep. 2016 Jun 27;6:28691. https://doi.org/10.1038/srep28691 PMID: 27345436; PMCID: PMC4922018.

35. Watanabe-Asaka T, Niihori M, Terada M, Oda S, Iwasaki K, Sudoh M, et al. Technology with High-Speed Movies to Analyze the Movement of Internal Organs in Medaka. Transac Jpn Soc Aeronautica Space Sci Aerospace Technol Jpn. 2012;10 (ists28):1–4. https://doi.org/10.2322/tastj.10.Pp_1

36. Watanabe-Asaka T, Sekiya Y, Wada H, Yasuda T, Okubo I, Oda S, et al. Regular heartbeat rhythm at the heartbeat initiation stage is essential for normal cardiogenesis at low temperature. BMC Dev Biol. 2014 Feb 25;14:12. https://doi.org/10.1186/1471-213X-14-12 PMID: 24564206; PMCID: PMC3936829.

37. Iwamatsu T. The integrated book for the biology of the medaka. Okayama: University Education Press; 1997

38. Kawasaki T, Saito K, Deguchi T, Fujimori K, Tadokoro M, Yuba S, et al. Pharmacological characterization of isoproterenol-treated medaka fish. Pharmacol Res. 2008 Nov-Dec;58(5-6):348–55. https://doi.org/10.1016/j.phrs.2008.09.011 PMID: 18951980.

39. Matsui K. Temperature and heart beat in a fish embryo, Oryzias latipes. Sci Rep Tokyo Bunrika Daigaku. 1941;B5:313–324.

40. Matsuura Y. Effect of temperature on heart beat in Oryzias latipes. Zool Mag. 1933;45:367–368.

41. Tsukuda H. Temperature accommodation of heart beats in fresh water teleosts. Zool Mag. 1962;71:46.

42. Kojima T, Neishi H, Yoshizaki Y, Soeda H. Power Spectrum Analysis of Fish Heart Rate Variability Using Maximum Entropy Method. Fisheries Eng. 2001;38(2):145–150. https://doi.org/10.18903/fisheng.38.2_145

43. Introna R, Yodlowski E, Pruett J, Montano N, Porta A, Crumrine R. Sympathovagal effects of spinal anesthesia assessed by heart rate variability analysis. Anesth Analg. 1995 Feb;80(2):315–321. https://doi.org/10.1097/00000539-199502000-00019 PMID: 7818119.

44. Jänig W. Spinal cord reflex organization of sympathetic systems. Prog Brain Res. 1996;107:43–77. https://doi.org/10.1016/s0079-6123(08)61858-0 PMID: 8782513.

45. Kumada M, Terui N, Kuwaki T. Arterial baroreceptor reflex: its central and peripheral neural mechanisms. Prog Neurobiol. 1990;35(5):331–361. https://doi.org/10.1016/0301-0082(90)90036-g PMID: 2263735.

46. William K. Milsom. Mechanisms of ventilation in lower vertebrates: adaptations to respiratory and nonrespiratory constraints. Can J Zool. 1989;67(12):2943–2955. https://doi.org/10.1139/z89-417

47. Randall DJ. The nervous control of cardiac activity in the tench (Tinca tinca) and the goldfish (Carassius auratus). Physiol Zool. 1966;39(3):185–192 https://doi.org/10.1086/physzool.39.3.30152846

